# Calaxin is essential for the transmission of Ca^2+^-dependent asymmetric waves in sperm flagella

**DOI:** 10.1101/2020.09.22.309153

**Authors:** Kogiku Shiba, Shoji A Baba, Eiji Fujiwara, Kazuo Inaba

## Abstract

Regulation of waveform asymmetry in sperm flagella is critical for changes in sperm swimming trajectory as seen during sperm chemotaxis towards eggs. Ca^2+^ is known as an important regulator of asymmetry in flagellar waveforms. A calcium sensor protein, calaxin, which is associated with the outer arm dynein, plays a key role in the sperm waveform regulation in a Ca^2+^-dependent manner. However, the molecular mechanism underlying the regulation of asymmetric waves by Ca^2+^ and calaxin remains unclear. We performed experiments using caged ATP to elucidate the formation and propagation of asymmetric flagellar waves in the sperm of the ascidian *Ciona intestinalis*. Demembranated sperm cells were suspended in a solution containing caged ATP and reactivated using UV flash photolysis. Initial bends were formed at the base and propagated towards the tip of flagella; however, the bend direction was different between asymmetric and symmetric waves. A calaxin inhibitor, repaglinide, had no effect on initial bend formation, but significantly inhibited the generation of the second flagellar bend in the reverse direction, resulting in the failure of asymmetric wave formation and propagation. These results suggest that calaxin plays a critical role in Ca^2+^-dependent transmission of flagellar asymmetric waveforms.

## INTRODUCTION

Changes in the bending pattern of cilia and flagella are important for the regulation of cell movements and extracellular fluid flow (Inaba, 2015). In particular, the sperm flagellar waveform is generally composed of two bends: a principal bend (P-bend) and a reverse bend (R-bend) (Goldstein, 1977, Brokaw, 1979, Gibbons and Gibbons, 1980, Inaba and Shiba, 2018). When the radii of curvature of two opposite flagellar bends are almost the same, sperm cells generate symmetric waveforms and swim straight. On the other hand, when the radius of curvature of the P-bend is much larger than that of the R-bend, sperm cells generate highly asymmetric waveforms, resulting in circular movements. The regulation of asymmetry in sperm flagellar waveforms is critical for the changes of sperm swimming direction, as seen during sperm chemotaxis towards the eggs (Miller, 1985, Wachten et al., 2017, Yoshida and Yoshida, 2011).

Ca^2+^ is known to play a key role in the regulation of asymmetry in flagellar waveforms. For instance, the flagellar waveforms in demembranated models of sea urchin sperm cells change from symmetric to asymmetric with increasing Ca^2+^ concentrations (Brokaw, 1979, Gibbons and Gibbons, 1980). Such a change is implied in the process of egg fertilisation, as Ca^2+^ bursts induce highly asymmetric waveforms during sperm chemotaxis, thereby inducing turning of sperm towards the egg in ascidians and sea urchins (Shiba et al., 2008, Guerrero et al., 2010, Bohmer et al., 2005, Wood et al., 2005). However, the Ca^2+^-dependent molecular mechanism regulating the formation and propagation of asymmetric waves remains unclear.

We previously identified a novel Ca^2+^-binding axonemal protein, calaxin, in the sperm of the ascidian *Ciona intestinalis* (recently renamed *C. intestinalis* type A or *Ciona robusta*). Calaxin is a member of the neuronal calcium sensor (NCS) family and directly interacts with outer arm dynein (Mizuno et al., 2009). Notably, inhibition of calaxin with a specific NCS inhibitor, repaglinide, suppresses the propagation of asymmetric flagellar waveforms required for chemotactic turn, although a transient increase in intracellular Ca^2+^ during the turning movement normally occurs (Mizuno et al., 2012). However, the mechanism by which calaxin contributes to the regulation of asymmetric wave formation remains largely unexplored. To fill this knowledge gap, we performed experiments using caged ATP to capture instantaneous images of the formation and propagation of asymmetric waves. Our results elucidated the mechanism by which asymmetric waves form and propagate, as well as the importance of calaxin for their regulation. These findings open new avenues towards a mechanistic understanding of flagellar motility, with possible applications in the fields of reproductive medicine and technology.

## RESULTS

To elucidate the mechanism underpinning the formation and propagation of asymmetric flagellar waves, we performed experiments using caged ATP to capture initial bend formation and subsequent propagation. In particular, demembranated *Ciona* sperm was incubated with 1 mM of caged ATP, and then reactivated using UV flash photolysis. This method allows the release of ATP from its caged compound by the UV flash within a few seconds (McCray et al., 1980, Goldman et al., 1982). To record the changes in sperm flagellar waveforms upon UV flashing, we used a phase-contrast microscope with a red LED for illumination and captured the images using a high-speed camera, to which the LED pulse was synchronised. UV flashes were applied through the objective lens by an UV LED located in the lamp house for observation of fluorescence.

First, we estimated the concentration of ATP released from caged ATP after UV flashing by measuring flagellar beat frequency. The beat frequency of demembranated sperm that had been reactivated with 1 mM of caged ATP and a 150 ms UV flash was estimated to be approximately 10 Hz (Fig. 1A). Notably, sperm beat frequency did not significantly differ between solutions with low (pCa10) and high (pCa5) Ca^2+^ concentrations. Further, we measured the beat frequency of demembranated and reactivated sperm at various concentrations of Mg-ATP (Fig. 1B). From the calibration curve, we estimated that ~0.1 mM of ATP was released under these conditions.

**Figure 1.**
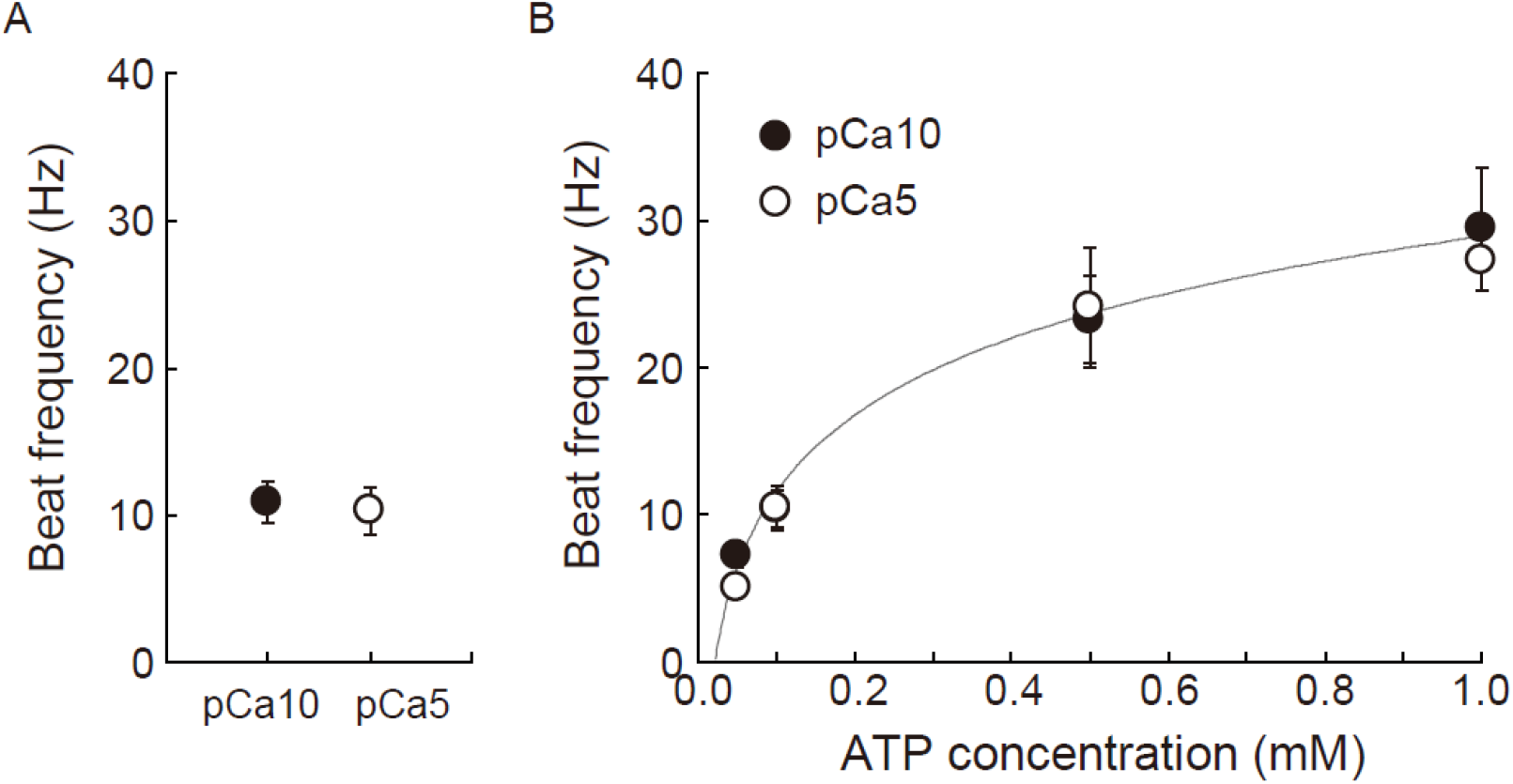
Estimation of the concentration of ATP released from caged ATP. A: Beat frequency of demembranated *Ciona* sperm incubated with 1 mM caged ATP and activated by a 150 ms UV flash. N = 11 (pCa10), and N = 12 (pCa5). B: Beat frequency of demembranated and reactivated *Ciona* sperm by various concentrations of Mg-ATP. Closed and open circles show Ca^2+^ concentrations in the reactivation solutions pCa10 and pCa5, respectively. N = 13–28 from three different experiments. Values are expressed as mean± S.D.

Next, we analysed the formation and propagation of flagellar waveforms upon ATP release. At a low Ca^2+^ concentration, released ATP induced the formation of an initial bend at the base of the flagellum 100–150 ms after the UV flash (Fig. 2A, Table 1, Movie S1). The bend propagated towards the flagellar tip, and the formation of a second bend was observed within 200 ms. Interestingly, low and high Ca^2+^ concentrations induced symmetric and asymmetric waveforms, respectively (Gibbons and Gibbons, 1980, Brokaw, 1979, Inaba and Shiba, 2018). Similar response times were observed in the formation and propagation of flagellar waveforms at high concentrations of Ca^2+^, although the curvature of the initial bend was larger than that of the second one (Fig. 2B, Table 1, Movie S2).

**Figure 2.**
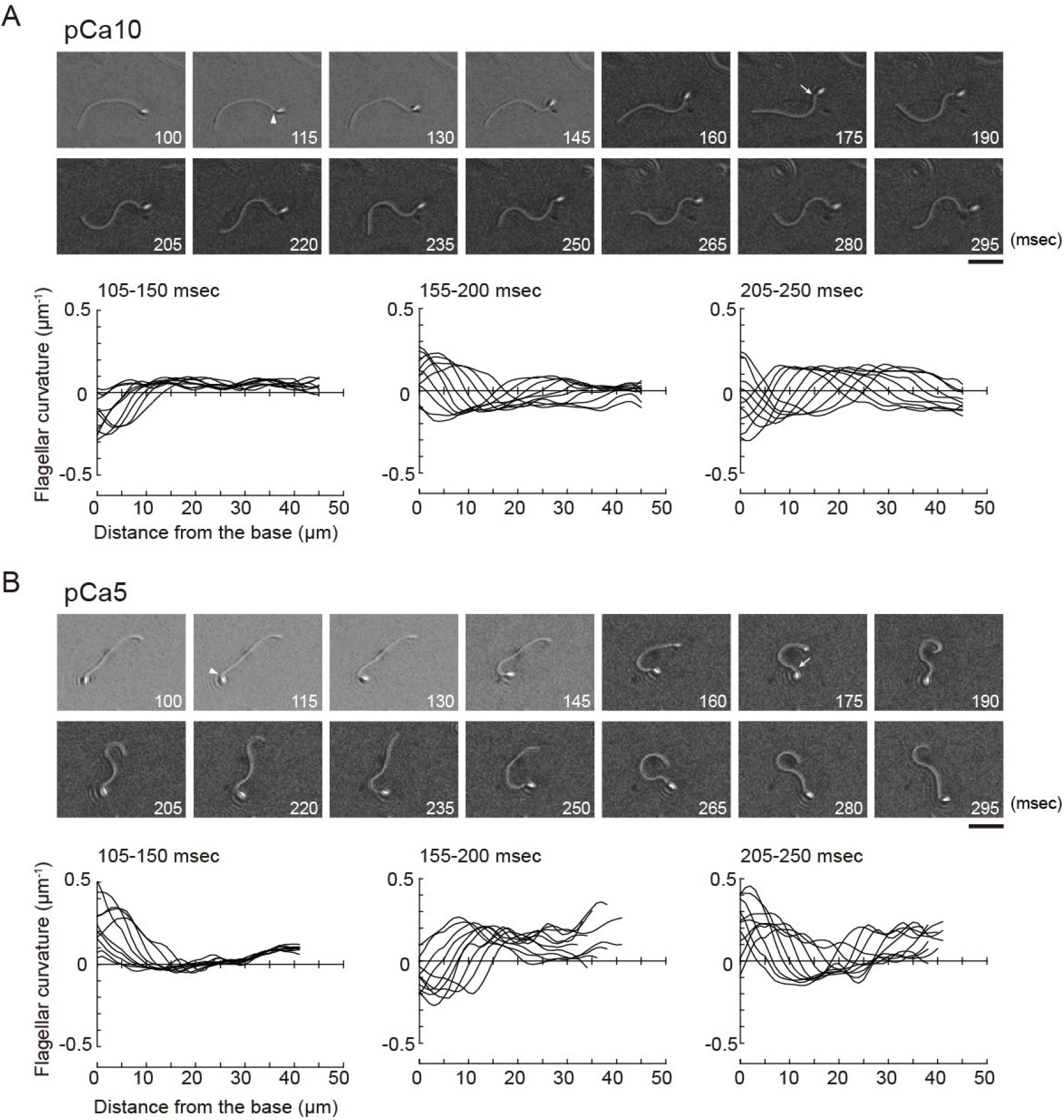
Generation of symmetric and asymmetric flagellar waveforms in *Ciona* sperm. Sperm cells were demembranated and reactivated through the photolysis of caged ATP in low (pCa10, A) or high (pCa5, B) Ca^2+^ concentrations. Upper panel: sequential images of sperm flagellar waveforms at 15 ms-intervals from 100 ms after the UV flash. Arrow heads and arrows indicate the initial bend and the second bend, respectively. Scale bar, 20 μm. Lower panel: changes of flagellar curvature occurring 105–250 ms after the UV flash are plotted against the distance from the base of flagellum. Ten waveforms produced at 50 ms are overwritten. Symmetric and asymmetric waveforms were generated in low calcium concentrations (pCa10; A) and high calcium concentrations (pCa5; B), respectively.

**Table 1.**
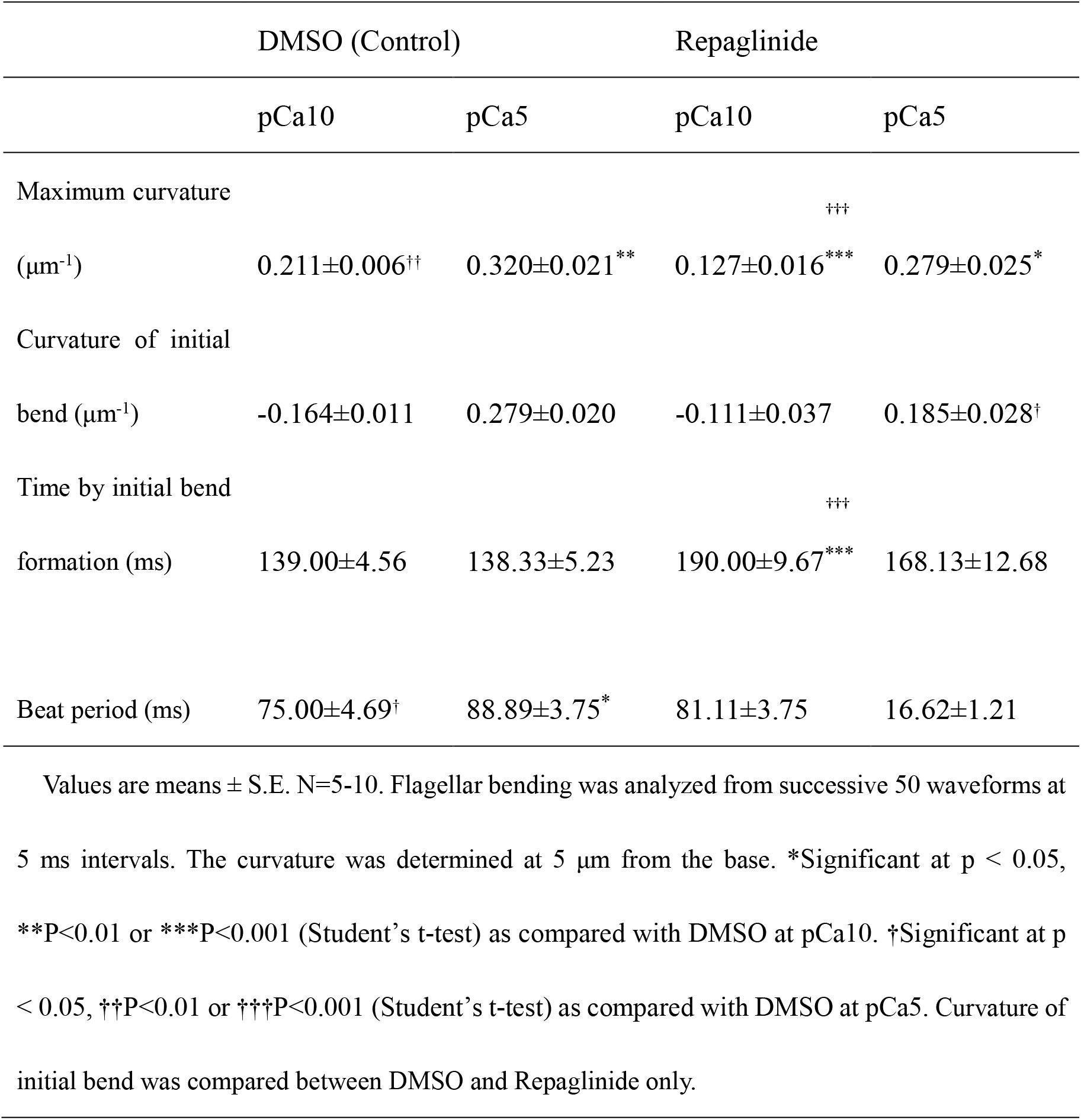
Sperm flagellar bending generated by the photolysis of caged ATP.

We previously reported that calaxin is an important factor regulating the propagation of asymmetric waveforms (Mizuno et al., 2012). Therefore, here we examined the role of calaxin in bend formation and propagation during initial generation of asymmetric flagellar waves through the use of a calaxin inhibitor, repaglinide. Repaglinide is an inhibitor of NCS family proteins (Okada et al., 2003), which specifically binds to calaxin and inhibits sperm chemotaxis in *Ciona* sperm flagella (Mizuno et al., 2012). Demembranated sperm cells were incubated with caged ATP and repaglinide and irradiated with UV light to induce ATP release. The process of initial bend formation and propagation was recorded at both low and high Ca^2+^ concentrations. At low concentrations of Ca^2+^, both the initial bend and the second bend were formed normally; however, such formation occurred a little later (~250 ms) upon repaglinide treatment when compared to that in the control (Fig. 3A, Table 1, Movie 3). Moreover, the maximum curvature after UV flashing was significantly smaller in the flagella of repaglinide-treated sperm than in the absence of repaglinide (Table 1). In addition, at a high concentration of Ca^2+^ in the presence of repaglinide, the initial bend was formed, but did not propagate to the flagellar tip (Fig. 3B, Movie S4). On the other hand, in the absence of repaglinide, the bend formed at the base and then increased its curvature, but the curvature at the posterior region remained quite limited. Moreover, the flagellum never formed the second bend, reflecting a quiescence pattern that was previously reported in sea urchin sperm (Goldstein, 1979; Gibbons and Gibbons, 1980). Therefore, the flagellum continued to vibrate at a high frequency with an overall constant waveform (Movie S4).

**Figure 3.**
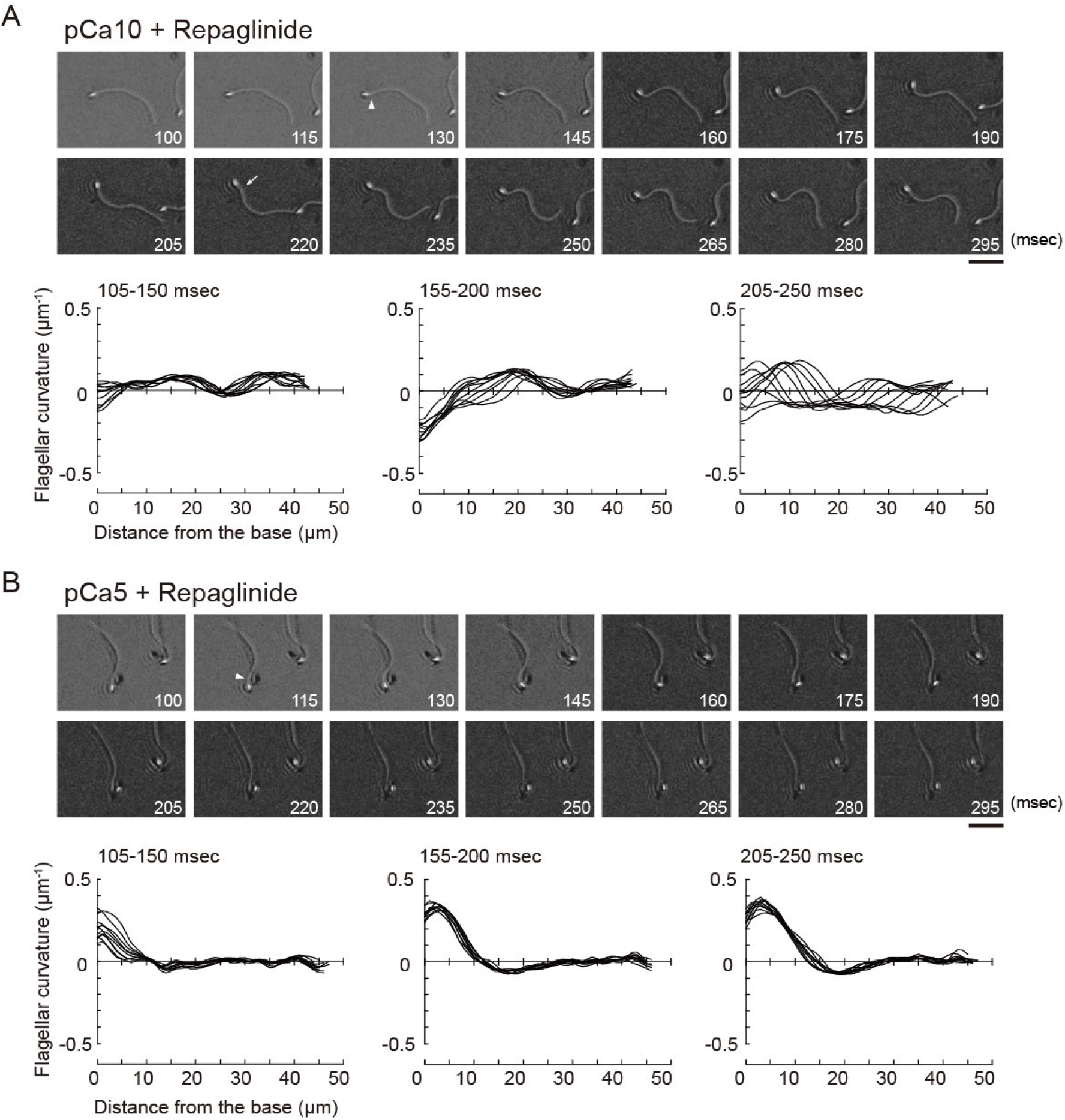
Effect of a calaxin inhibitor on the propagation of asymmetry waveforms at high calcium concentrations. Sperm cells were demembranated and reactivated through the photolysis of caged ATP in low (pCa10, A) or high (pCa5, B) Ca^2+^ concentrations with 150 μM repaglinide. Upper panel: sequential images of sperm flagellar waveforms at 15 ms-intervals from 100 ms after the UV flash. Arrow heads and arrows indicate the initial bend and the second bend, respectively. Scale bar, 20 μm. Lower panel: changes of flagellar curvature occurring 105–250 ms after the UV flash are plotted against the distance from the base of flagellum. Ten waveforms produced at 50 ms are overwritten.

Interestingly, we found that the initial bends of sperm reactivated at low Ca^2+^ concentrations consisted mostly of R-bends (Fig. 2A). In contrast, the initial bends formed in the sperm reactivated at high Ca^2+^ concentrations were almost exclusively P-bends (Fig. 2B). The same pattern was also observed in the presence of repaglinide (Fig. 3A and 3B). Moreover, quantitative analysis showed that ~80–90% of the sperm flagella formed an initial bend in this manner (Fig. 4), suggesting that the direction of the initial bend depends on Ca^2+^.

**Figure 4.**
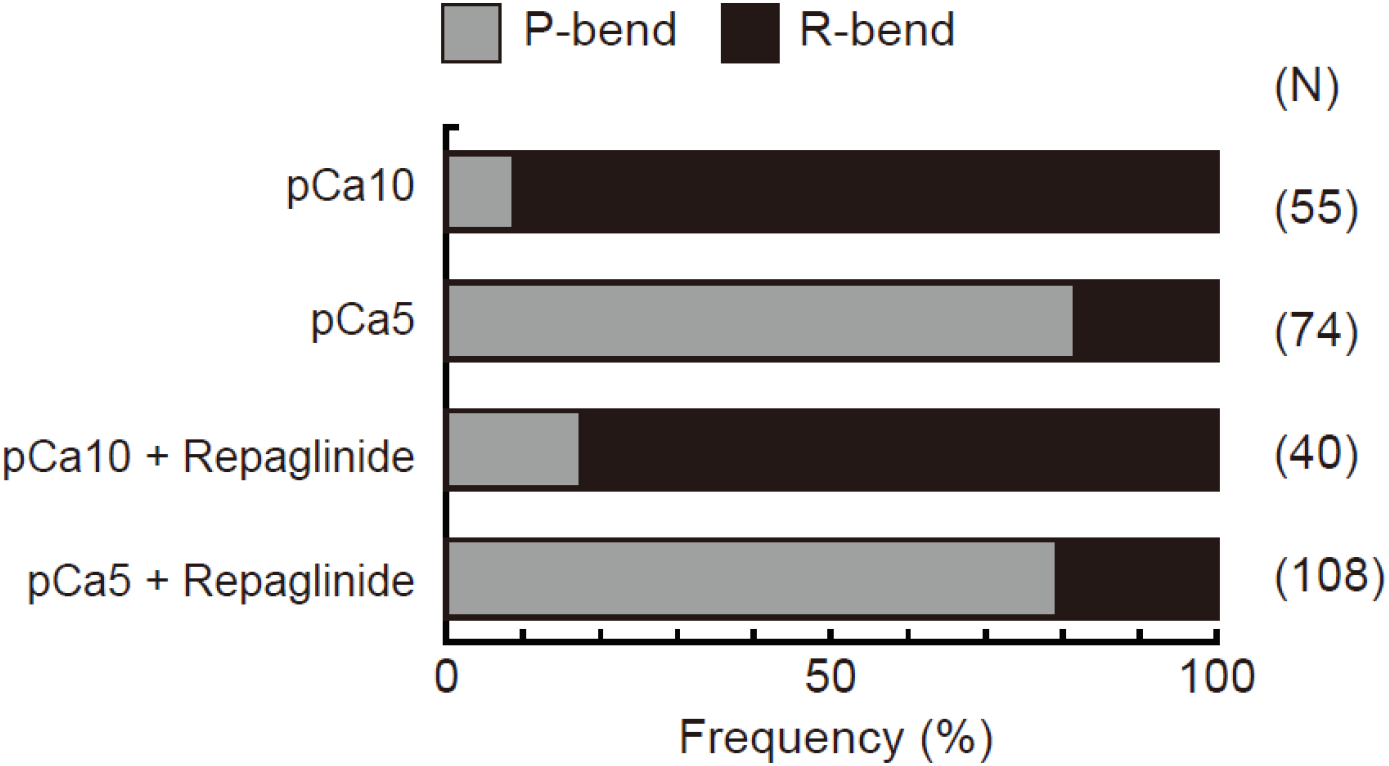
Role of the preferential direction of the initial bend in the generation of symmetric and asymmetric flagellar waveforms. The relative frequencies of principal (P) and reverse (R) bends are shown for the initial bends formed after activation of demembranated *Ciona* sperm. Sperm was reactivated by photolysis of caged ATP in low (pCa10) or high (pCa5) Ca^2+^ concentrations in the presence of 0.5% DMSO (control) or 150 μM repaglinide. In both control and repaglinide-treated sperm, the formation of flagellar bends started with R-bends at pCa10, whereas it started with P-bends at pCa5. The number in brackets represents the number of observed spermatozoa in three different experiments.

For a better understanding of the process of flagellar bend formation, the changes in flagellar curvature were plotted against time and distance from the flagellar base in a three-dimensional (3D) graph (Fig. 5). At low Ca^2+^ concentrations, R-bends were generated at the flagellar base and propagated to the tip, followed by generation and propagation of P-bends (Fig. 5A). On the other hand, P-bend formation preceded R-bend generation at high Ca^2+^ concentrations (Fig. 5B). Repaglinide had little effect on the overall pattern of bend formation at low Ca^2+^ concentrations, although the propagation of R-bend proceeded more slowly and, consequently, the formation of the second bend (P-bend) was delayed (Fig. 5C, Table 1). Conversely, at high Ca^2+^ concentrations, repaglinide did not inhibit the generation of P-bends at the flagellar base, but suppressed both their propagation and the subsequent formation of R-bends. In these conditions, P-bends were formed completely ~150 ms after UV flashing and maintained a constant curvature for up to 325 ms (Fig. 5D). When observing the vibration of the flagellum (Movie S4), the curvature of the P-bend oscillated with a periodicity of 5–40 ms and a frequency of 75.14 ± 7.49 Hz (N = 6) (Asterisks in Fig. 5D; Table 1).

**Figure 5.**
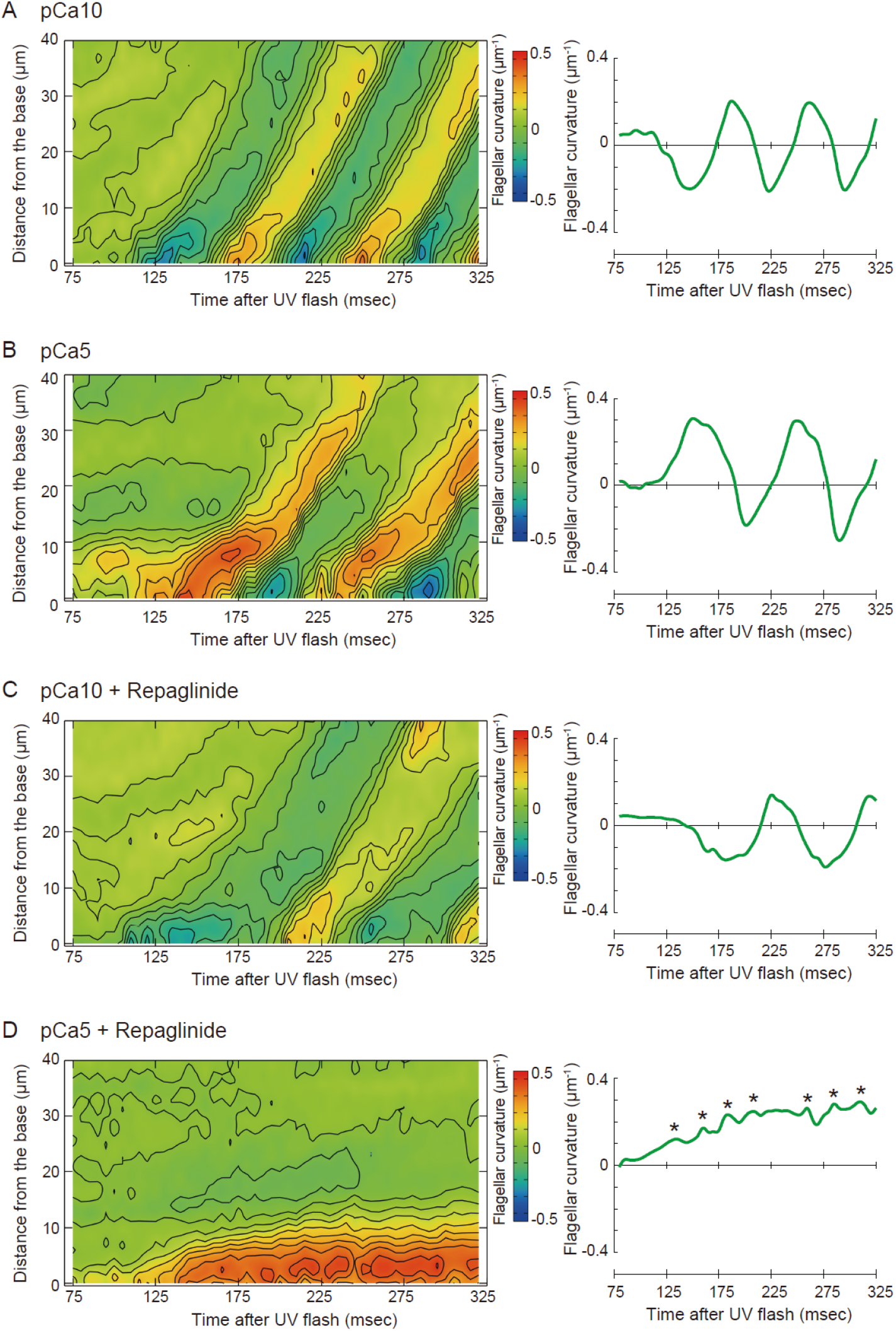
Pseudocolor maps showing spatiotemporal changes of the flagellar curvature. Left panels: the flagellar curvature of demembranated *Ciona* sperm was plotted against the distance from the base and time after the UV flash by pseudocolor mapping. Right panels: the flagellar curvature at 5 μm from the base was plotted against time after the UV flash. Sperm was reactivated through photolysis of caged ATP in low (pCa10; A, C) or high (pCa5; B, D) Ca^2+^ concentrations in the presence of 0.5% DMSO (control; A, B) or 150 μM repaglinide (C, D).

## DISCUSSION

Periodical flagellar oscillation is regulated by a highly sophisticated machinery composed of dynein motors, microtubules, and other regulatory factors (Inaba, 2003). In particular, local sliding between doublet microtubules by dyneins is essential for bend formation (Shingyoji et al., 1977). Indeed, coordinated activation or inactivation of dyneins within nine sets of doublet microtubules along the flagella controls the bend curvature, bending direction, wavelength, and propagation velocity (Inaba, 2011, Lin and Nicastro, 2018, King, 2018). For instance, switching of dynein activity between doublet 7 and 3 is thought to be responsible for the periodical and planar oscillation of sperm flagella. Although this switching is considered to be precisely regulated by chemical and mechanical signals, the details of this regulatory mechanism remain unclear.

In this study we developed a microscopic illumination system using LED light and applied it to examine the initiation and subsequent propagation of flagellar bends induced by caged ATP. This experimental system is very useful for observing the formation and propagation of flagellar waveforms without physical disturbances such as fluid flow and mechanical stimuli (Vernon and Woolley, 1994, Tani and Kamimura, 1998, Hayashi and Shingyoji, 2008). Moreover, photolysis of caged ATP can induce the sliding of microtubules at the desired time. Subsequently, the mechanical load at the base of the flagellum converts such sliding into the formation of the first bend at the proximal region. Thus, our system allows to capture the moment of first bend formation.

With this experimental setting, we found that the first bend of symmetric or asymmetric waves formed at the proximal region of the flagellum is predominantly an R-bend or a P-bend, respectively (Figs 4 and 5). Studies in sea urchins showed that the formation of P-bends and R-bends is induced by the activation of dyneins on doublet 7 and 3, respectively (Sale, 1986, Shingyoji, 2018). Therefore, the initiation of the first R-bend in a symmetric flagellar wave under low Ca^2+^ conditions in *Ciona* (Fig. 2A) might be induced by the activation of dyneins on doublet 3 as well. Moreover, the formation of a second P-bend at the base of the flagellum after propagation of the R-bend towards the flagellar tip might depend on the switch of active dyneins from doublet 3 to doublet 7 (Fig. 6, left).

**Figure 6.**
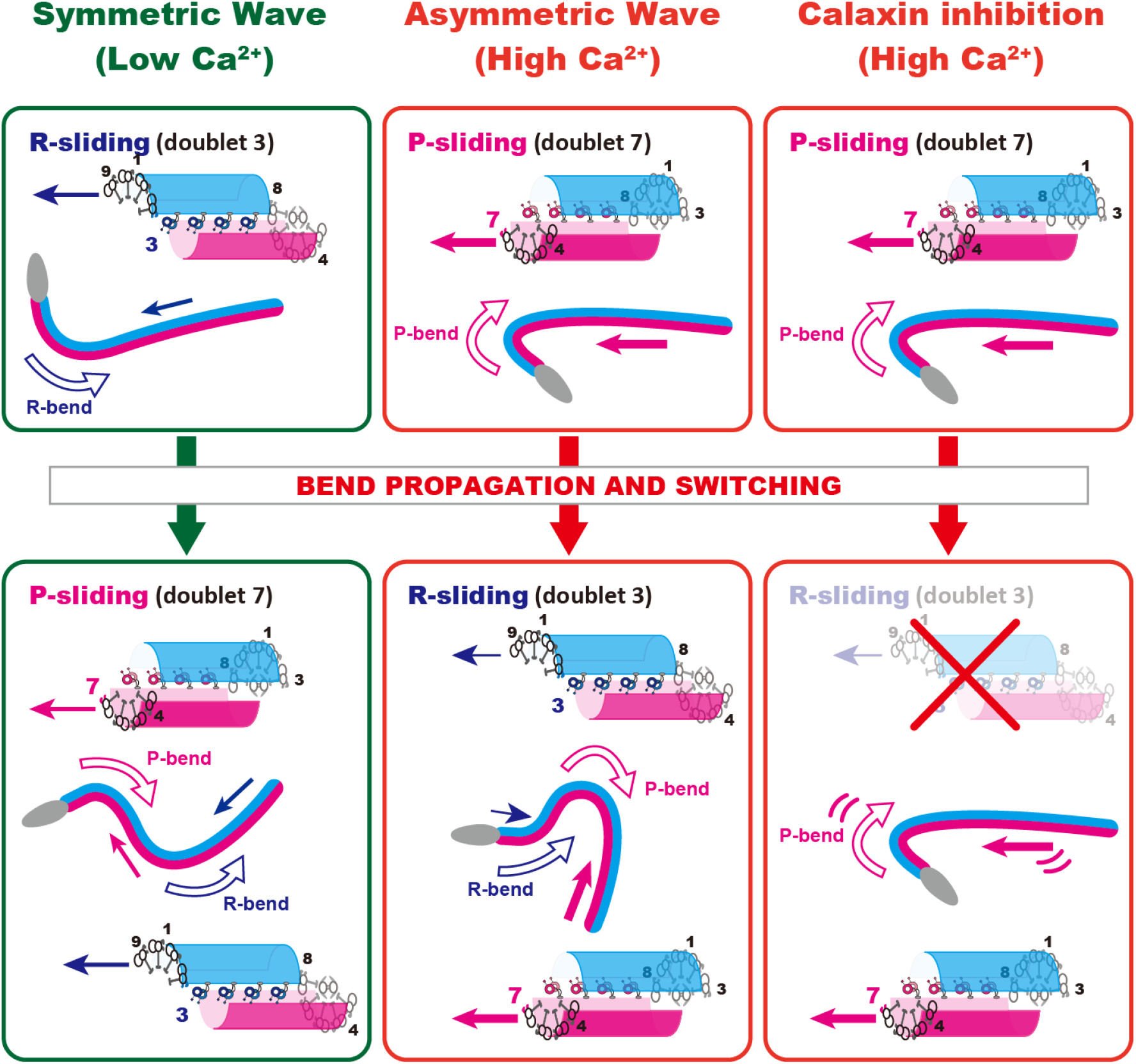
A model for calaxin-dependent transmission of asymmetric flagellar waves. The putative mechanisms mediating initial bend formation and its propagation are shown under different experimental conditions. Left: Formation of a symmetric wave under low Ca^2+^ conditions. The first R-bend is formed by microtubule sliding induced by doublet 3 dynein. In turn, this R-bend induces switching of active dynein from doublet 3 to 7, resulting in microtubule sliding induced by doublet 7 dynein to form a P-bend. Middle: Generation of an asymmetric wave under high Ca^2+^ conditions. A P-bend is first formed by microtubule sliding induced by doublet 7 dynein. In turn, this P-bend induces the switching of active dynein from doublet 7 to 3. However, the activity of doublet 3 dynein is suppressed by calaxin, resulting in the formation of an R-bend with lower curvature and in the subsequent production of an asymmetric wave. Dynein-driven microtubule sliding is known to be suppressed by calaxin at high Ca^2+^ concentrations; therefore, it is plausible that a mechanism for the relief of calaxin-mediated inhibition on doublet 7 dynein is involved in P-bend formation (see text). Right: Effects of the calaxin inhibitor repaglinide on asymmetric wave formation under high Ca^2+^ conditions. An initial P-bend is normally formed. However, this does not propagate or trigger the formation of an R-bend. Repaglinide releases calaxin-dependent inhibition of microtubule sliding in the R-bend, but does not affect other putative calaxin-independent mechanisms for the regulation of microtubule sliding, such as the transmission of a feedback mechanical load from the P-bend.

On the other hand, high Ca^2+^ specifically suppresses the activation of doublet 3 dyneins and attenuates the formation of R-bends in sea urchin sperm flagella (Nakano et al., 2003). Similarly, in *Ciona* sperm, bend initiation occurs with the formation of a P-bend at the proximal region of the flagellum under high Ca^2+^ conditions, followed by the propagation of such P-bend and the formation of an R-bend at the flagellar base (Figs 2 and 4). Therefore, we suggest that, following Ca^2+^ bursts, the activation of dyneins occur predominantly on doublet 7, thereby inducing an initial P-bend at the proximal region of the flagellum. Subsequent formation of the small R-bend near the base of the flagellum would follow the activation of dyneins on doublet 3. On the other hand, the sliding of doublet 3 (R-bend) would be suppressed under high Ca^2+^ conditions, resulting in the formation of an asymmetric wave (Fig. 6, middle).

Moreover, the specific role of the Ca^2+^-binding protein calaxin in this mechanism could be dissected through the use of the NCS inhibitor repaglinide. In the presence of repaglinide, the initial P-bend was normally formed under high Ca^2+^ conditions. However, it did not propagate and, as a consequence, the R-bend never appeared (Fig. 3B and 5D). Therefore, calaxin is probably essential for the activation of doublet 3 dyneins to form R-bends, but not for the activation of doublet 7 dyneins to form P-bends (Fig. 6, right). Considering also the results relative to low Ca^2+^ concentrations, we propose that calaxin is primarily involved in the regulation of R-bend formation and propagation.

To complete the model depicted in Fig. 6, three key questions need to be addressed. First, the mechanism by which R-bend is initially formed by the sliding of doublet 3 at the proximal region of the flagellum in low Ca^2+^ conditions should be investigated. Indeed, flagellar bends are formed through the conversion of microtubule sliding into bending of physical structures by mechanical resistance (Shingyoji et al., 1977). Therefore, the preference of doublet 3 sliding during the initial steps of bend formation could be due to the difference in mechanical sensitivity between the sliding of doublets 3 and 7. In fact, the preference for R-bends during bend initiation has also been shown to occur during flagellar beating induced by head vibration in sea urchin sperm (Eshel et al., 1991). It is possible that asymmetric structures in the axoneme, such as the central pair apparatus (CP) / radial spokes (RS), determine such an initial preference.

Secondly, it would be interesting to know if the P-bend can be formed under high Ca^2+^ conditions (Fig. 6, middle). We previously demonstrated by an *in vitro* motility assay using purified microtubules and outer arm dynein that dynein-mediated microtubule sliding was suppressed by calaxin under high Ca^2+^ conditions (Mizuno et al., 2012). Since calaxin is associated with outer arm dyneins on all nine doublet microtubules (Mizuno et al., 2009), there must be a mechanism mediating the relief of calaxin-dependent inhibition of doublet 7 sliding. It is possible that the activity of doublet 7 dynein is regulated by CP/RS, which possibly relieves the Ca^2+^/calaxin-dependent inhibition of outer arm dynein through protein phosphorylation, similar to their known role in the modulation of the activity of inner arm dynein (Habermacher and Sale, 1997, King and Dutcher, 1997, Smith and Yang, 2004). Therefore, the activation of doublet 7 dynein under high Ca^2+^ conditions might occur by a calaxin-independent mechanism. However, other calcium-binding proteins, calmodulin and centrin, are known to be localised in these structures (Yang et al., 2001, Dymek and Smith, 2007) as well as in inner arm dyneins (Piperno et al., 1992, Yamamoto et al., 2008, Yagi et al., 2009).

Thirdly, it is worth investigating the mechanism by which repaglinide inhibits both the formation and the propagation of R-bends under high Ca^2+^ conditions (Fig. 6, right). We previously showed that repaglinide rescues calaxin-dependent inhibition of microtubule sliding in *in vitro* motility assays (Mizuno et al., 2012). These results imply the existence of a mechanism for the inhibition of microtubule sliding on doublet 3 dyneins driving the formation and propagation of R-bends. Although this has not yet been elucidated, we speculate that the formation of P-bends exerts a mechanical load on the proximal region, where calaxin is essential for the activation of doublet 3 dyneins and the induction of microtubule sliding, triggering the formation of R-bends under the loaded state. Repaglinide would inhibit this activity of calaxin, resulting in the suppression of R-bend formation and propagation (Fig. 6, right).

In conclusion, the formation and propagation of asymmetric waves in sperm flagella depend on Ca^2+^ concentration and calaxin activity, although the detailed mechanism is yet to be elucidated. Possibly, Ca^2+^/calaxin-dependent regulation of dynein activity might play a key role in determining the unequal activity of microtubule sliding between the two sides of the flagellum. Further studies on the role of calaxin in the regulation of outer arm dyneins by mechanical loading or phosphorylation will shed light on the oscillatory mechanism of flagellar and ciliary movement.

## MATERIAL AND METHODS

### Materials

The ascidian *C. intestinalis* (type A; also called *C. robusta*) was collected from Onagawa Bay near the Onagawa Field Research Center, Tohoku University or obtained from the National BioResource Project for *Ciona* (http://marinebio.nbrp.jp/). Animals were kept in aquaria under constant light for accumulation of gametes without spontaneous spawning. Semen samples were collected by dissecting the sperm duct and kept on ice until use.

### Chemical and solutions

Ca^2+^-free sea water (CFSW) contained 478.2 mM NaCl, 9.39 mM KCl, 48.27 mM MgCl_2_, 2.5 mM EGTA and 10 mM Hepes-NaOH (pH 8.0). Sperm demembranation solution contained 0.2 M potassium acetate, 1 mM MgSO_4_, 2.5 mM EGTA, 20 mM Tris-acetate (pH 7.8), 0.04% Triton X-100 and 1 mM DTT. Pre-incubation buffer contained 0.15 M potassium acetate, 2 mM MgSO_4_, 2.5 mM EGTA, 39 μM (pCa10) or 2.51 mM (pCa5) CaCl_2_, 50 mM Tris-HCl (pH8.0) and 1 mM DTT. Reactivation solution was the same as pre-incubation buffer but with 2 mM ATP. Free Ca^2+^ concentration was assessed by CALCON (http://www.bio.chuo-u.ac.jp/nano/calcon.html). Caged ATP was purchased from Dojindo (Kumamoto, Japan) dissolved in Pre-incubation buffer. Repaglinide was purchased from Sigma-Aldrich (St Louis, MO) and dissolved in dimethyl sulfoxide (DMSO). Theophylline from Sigma-Aldrich was dissolved in CFSW. Other reagents were of analytical grades.

### Preparation of demembranated sperm

Demembranation and reactivation of *C. intestinalis* sperm were performed as described previously with some modifications (Mizuno et al., 2012). Semen was suspended in 100 volumes of CFSW containing 1 mM theophylline in CFSW to activate motility and incubated for 5 min at 25°C. Sperm were demembranated with 10 volumes of demembranation solution incubated for 3 min at 25°C. Demembranated sperm were kept for several minutes in pre-incubation buffer with or without 1 mM caged ATP until use. Demembranated sperm incubated with caged ATP were reactivated by UV irradiation. Demembranated sperm without caged ATP were reactivated with the same volume of the reactivation buffer containing 2 mM ATP (Final ATP concentration was 1 mM). DMSO (control) or Repaglinide was added to the demembranation solution, pre-incubation buffer and reactivation buffer. DMSO concentration was kept below 0.5% in all experiments.

### Experimental system

Movements of demembranated sperm incubated with caged ATP were observed under an inverted microscope with phase optics (IX71, Olympus, Tokyo, Japan) with a 20x objective and recorded with a high-speed CCD camera (HAS220, Detect, Tokyo, Japan). Flagellar waveforms were captured by a red-LED (Edison 3WStar, Taipei, Taiwan) and a laboratory-made LED stroboscopic illumination system synchronized with the exposure signals from the high-speed camera. Images were taken at a frame rate of 200 fps with 0.2 msec pulse from the red LED. A UV-LED (NS365L-3SVR, 365 nm, Nitride Semiconductors, Tokushima, Japan) was used for the photolysis of caged ATP. A mercury lamp in the lamp house of the microscope was replaced by a laboratory-made LED holder with UV-LED. A laboratory-made LED controller generated a 150 msec pulse for UV irradiation and triggered a recording cue signal on the high-speed camera (HAS220) which enables to record images 0.25 sec before and 2.25 sec after UV irradiation. To measure the flagellar beat frequency of reactivated sperm with Mg-ATP flagellar waveforms were observed under an upright microscope with phase optics (BX51, Olympus) with a 20x objective and recorded with a high-speed CCD camera (HAS-D3, Detect) at 500 fps.

### Analysis of flagellar waveforms

The flagellar beat frequency and the flagellar curvature were analyzed using Bohboh software (Bohbohsoft, Tokyo, Japan). Individual images of sperm flagella were tracked automatically, and their curvatures calculated based on the method of Baba and Mogami (1985). Pseudocolor-maps of the flagellar curvature along the distance and time were created using by gnuplot 5.0 (http://www.gnuplot.info/).

### Statistical Analysis

All experiments were repeated at least three times with three different specimens. Data is expressed as means ± SE. Statistical significance was calculated using Student’s t-test; P < 0.05 was considered significant.

## ACKNOWLEDGEMENTS

We thank Drs Satoe Aratake, Reiko Yoshida, Manabu Yoshida, Yutaka Satou and other staff at Misaki Marine Biological Station, Grad Schl of Sci, University of Tokyo and in Lab of Dev Genomics, Grad Schl of Sci, Kyoto University, who distribute *Ciona* under the National BioResource Project, AMED, Japan. We are grateful to all the staff members of Onagawa Field Center, Graduate School of Agricultural Science, Tohoku University and International Coastal Research Center, University of Tokyo for supplying *Ciona*. This work was supported in part by grants from MEXT (Ministry of Education, Culture, Sports, Science and Technology), Japan for Innovative Areas (No. 15H01201) and Scientific Research (B) (No. 22370023) to K.I. and for Scientific Research (C) (No. 19K06592, No. 16K07337) K.S. and by JST-BIRD (Japan Science and Technology Agency-Institute for Bioinformatics Research and Development), Japan to K.I.

## Supplementary information

**Movie S1:** Generation of symmetric flagellar waveforms in *Ciona* sperm. Sperm cells were demembranated and reactivated through the photolysis of caged ATP in low (pCa10) Ca^2+^ concentrations with 0.5% DMSO. The video is processed by background subtraction method and plays at a speed of 0.05x. Scale bar; 20 μm. The character “UV” appears during UV irradiation.

**Movie S2:** Generation of asymmetric flagellar waveforms in *Ciona* sperm. Sperm cells were demembranated and reactivated through the photolysis of caged ATP in high (pCa5) Ca^2+^ concentrations with 0.5% DMSO. The video is processed by background subtraction method and plays at a speed of 0.05x. The character “UV” appears during UV irradiation.

**Movie S3:** Generation of symmetric flagellar waveforms in *Ciona* sperm. Sperm cells were demembranated and reactivated through the photolysis of caged ATP in low (pCa10) Ca^2+^ concentrations with 150 μM repaglinide. The video is processed by background subtraction method and plays at a speed of 0.05x. The character “UV” appears during UV irradiation.

**Movie S4:** Generation of asymmetric flagellar waveforms in *Ciona* sperm is inhibited by calaxin blocker. Sperm cells were demembranated and reactivated through the photolysis of caged ATP in high (pCa5) Ca^2+^ concentrations with 150 μM repaglinide. The video plays at a speed of 0.05x. Scale bar; 20 μm. The character “UV” appears during UV irradiation.

